# On the Design of a PID Bio-controller with Set Point Weighting and Filtered Derivative Action

**DOI:** 10.1101/2022.03.21.485223

**Authors:** Emmanouil Alexis, Luca Cardelli, Antonis Papachristodoulou

## Abstract

Effective and robust regulation of biomolecular processes is crucial for designing reliable synthetic bio-devices functioning in uncertain and constantly changing biological environments. Proportional-Integral-Derivative (PID) controllers are undeniably the most common way of implementing feed-back control in modern technological applications. Here, we introduce a highly tunable PID bio-controller with *set point weighting* and filtered derivative action presented as a chemical reaction network with mass action kinetics. To demonstrate its effectiveness, we apply our PID scheme on a simple biological process of two mutually activated species, one of which is assumed to be the output of interest. To highlight its performance advantages we compare it to PI regulation using numerical simulations in both the deterministic and stochastic setting.

## I. INTRODUCTION

The fast growing field of Synthetic Biology aims to engineer biomolecular systems with novel and useful functionalities in order to tackle a long list of pressing, real-world problems [1], [2], [3], [4]. Arguably, one of the main challenges of building synthetic bio-devices operating in the uncertain cellular environment is achieving a reliable and predictable behaviour. Feedback control theory provides a large variety of tools that have proven to be of fundamental importance in regulating such devices, optimizing their function and rendering them robust to disturbances [5], [6], [7], [8], [9].

Proportional - Integral - Derivative (PID) feedback controllers are regarded as the workhorses of control engineering [10], [11]. They are often called “three - term” controllers due to their triple control action accounting for the past, present and future. More specifically, integral control (I-term) accounts for the history of the error between the set point (desired target value) and the output of interest by accumulating it over time. An important characteristic of the I-term is its ability to eliminate the steady-state error, provided that the feedback system is stable. The present is represented by the P-term which produces a control signal proportional to the current value of the error. Lastly, derivative control (D-term) provides anticipatory action by estimating future values of the error via linear extrapolation.

Because of the pervasiveness of PID control in technological applications, the biomolecular implementation of PID controllers has been of great interest in Synthetic Biology and there have been several successful research efforts towards this direction. Notably, the authors in [12] present a hierarchical library of nonlinear PID controllers consisting of up to four biomolecular species with a first-order low-pass filter accompanying some or all the three control terms (P-, I- and D-term). Another example is the PID architecture proposed in [13] where different variations of Michaelis-Menten functions are exploited. Furthermore, the PID designs studied in [14], [15] use the so-called dual rail encoding [16], by which a signal is decomposed into two non-negative components and, thus, both positive and negative signals can be represented via biomolecular species. In addition, the authors [17] analyze the noise suppression properties of individual proportional, integral and derivative controllers tailored to gene expression.

In this paper, we introduce an alternative biomolecular network functioning as a PID controller around the nominal operation of the resulting closed-loop system (equilibria). The biomolecular interactions involved are defined by general chemical reaction networks (CRNs) based purely on mass action kinetics [18] and without using dual rail encoding. At the same time, our bio-controller acts solely on the target species (output of interest) without considering other species or reactions of the network to be controlled (open-loop system) as happens, for instance, in [12]. To achieve enhanced dynamic performance we adopt a special form of *set point weighting* commonly used in technological applications and we accompany derivative control with the strong filtering action of a second-order low-pass filter. Moreover, our PID configuration includes six controller species that allow us to build each of the P-, I-, D-terms almost independently providing significant tuning flexibility regarding controller gains, *set point weights* and filtering. Finally, the proposed PID configuration can be used for controlling any open-loop biological process assuming the existence of a biologically meaningful equilibrium and asymptotic stability for the resulting closed-loop system. Here we focus exclusively on scenarios where this condition is satisfied.

The paper is structured as follows: Section II presents some background concepts on PID control, biomolecular interactions, modelling tools and essential biomolecular motifs. Section III analyzes the main characteristics of the proposed PID bio-controller. Subsequently, an application example including a comparison between PI and PID control in the deterministic and stochastic setting is provided in Sections IV and V, respectively. Section VI concludes our work and discusses future research directions.

## II. BACKGROUND

In this section we first outline key properties of PID control and challenges in its implementation, review principles of biomolecular modelling and then present two important biomolecular motifs that implement integral and derivative action.

### A. Key points on PID control

Here we briefly present some important features of PID control action [10], [11] based on which our PID bio-controller has been developed.

First, recall that the “traditional”, ideal PID algorithm (Figure 1) is described as follows:

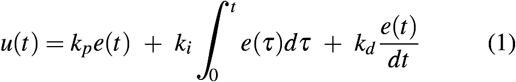

where *u*(*t*), *e*(*t*) represent the control input signal and the control error, respectively. The latter is defined as *e*(*t*) = *y_sp_* − *y*(*t*), where *y_sp_* is the set point and *y* is the process output.

**Fig. 1:**
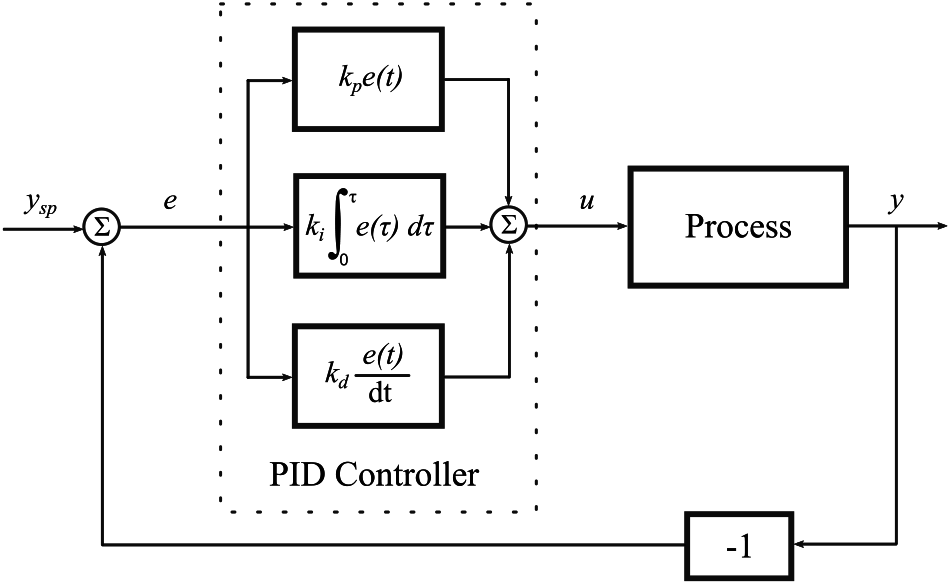
Ideal PID control of a process based on error feedback [10], [11].

A major problem of the (ideal) derivative action in Equation (1) is its sensitivity to high-frequency signal components. This can lead to excessively high gains and, by extension, large variations in terms of the control signal. A common strategy to overcome this obstacle is to accompany the derivative term with a low-pass filter.

Another challenge is *derivative kick*: When the set point is constant, the derivative of the error in Equation (1) becomes 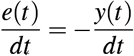 since 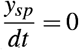. Abrupt changes of the set point (when the set point is adjusted) make the aforementioned derivative very large causing undesirable transients in the control signal (derivative kick). To avoid this, we can replace *e*(*t*) with −*y*(*t*) in the derivative term of Equation (1).

The behaviour of the controller can be further improved by modifying appropriately the error quantity on which the proportional action acts. To this end, we consider an alternative PID control law with *set point weighting*:

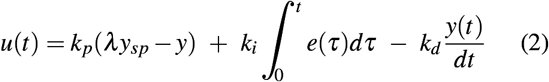

A PID controller based on Equation (2) with *λ* = 1 and *λ* = 0 is often referred to as a PI-D and I-PD controller, respectively. Finally, the error quantity in the integral term needs to remain unchanged in order for the error to go to zero at steady-state.

### B. Biomolecular interactions and modelling

In Figure 2A we present all different types of biomolecular interactions as well as their graphical notation used in this paper. These interactions can be divided into two main categories: non-catalytic reactions where the reactants are consumed in order for products to be formed and catalytic ones where species facilitate production/inhibition processes without being consumed.

**Fig. 2:**
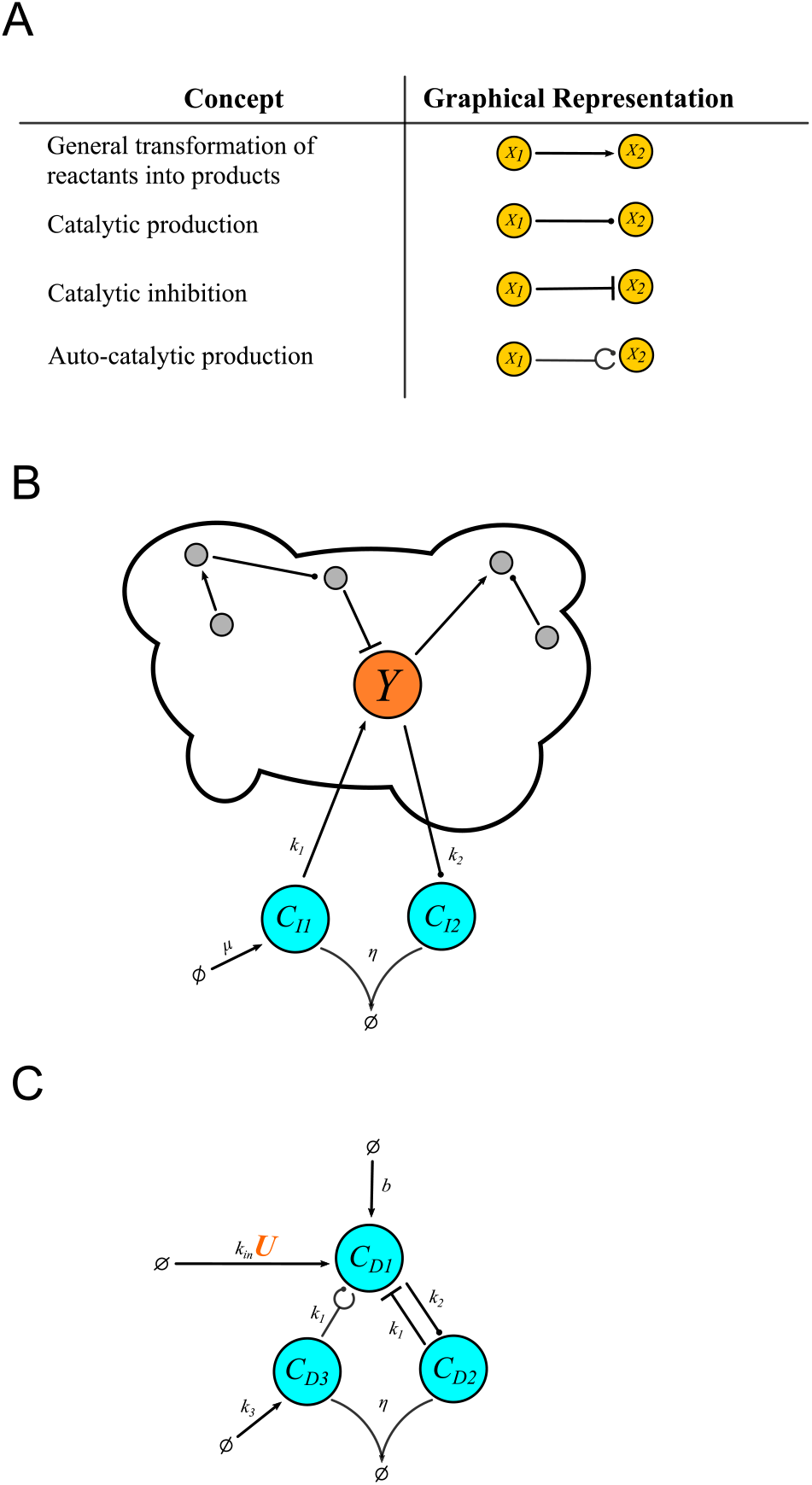
(A) Table with the different types of biomolecular interactions adopted from our previous work [19]. (B) *Antithetic* integral controller regulating a target species which is part of an arbitrary biological process - “cloud” network (CRN (3)). (C) *BioSD-III* differentiator module (CRN (5)).

For deterministic analysis of the biomolecular networks discussed in this paper (Sections II-C, III, IV) Ordinary Differential Equations (ODEs) following the law of mass action are used [18]. For the stochastic part (Section V) the Linear Noise Approximation (LNA) of the Chemical Master Equation (CME) [20], [21] is exploited.

### C. Two important biomolecular motifs

We now review the behaviour of two basic biomolecular motifs that were introduced in the literature, which are constituent elements of our PID architecture under appropriate modifications.

Figure 2B shows the *antithetic motif* introduced in [22], which is realized by controller species *C*_*I*1_, *C*_*I*2_, regulating a target (output) species, *Y*, which is part of an arbitrary biological process - “cloud” network. This mechanism can achieve *robust perfect adaptation* (RPA) through integral feedback control. To see this, focusing on the controller species, we have the CRN:

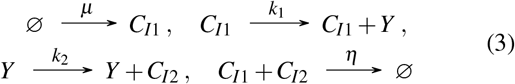

which can be modelled by the following set of ODEs:

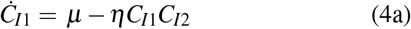

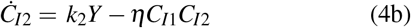

where *μ*, *k*_1_, *k*_2_, *η* ∈ ℝ_+_.

Integration is carried out by a hidden non physical “memory” variable. To see this, we subtract Equations (4a) - (4b) and integrate to obtain:

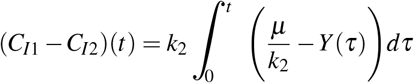

Consequently, assuming closed-loop stability, at the steady state:

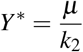

where the ^∗^ notation denotes the steady state of a variable.

Figure 2C shows a topology known as *BioSD-III* which we introduced in [19]. This topology is able to function as a signal differentiator module around its nominal operation. In particular, it receives an input signal, *U*, and calculates its filtered derivative in the output, in this case species *C_D_*_1_. To see this, the CRN for *BioSD-III* consists of the reactions:

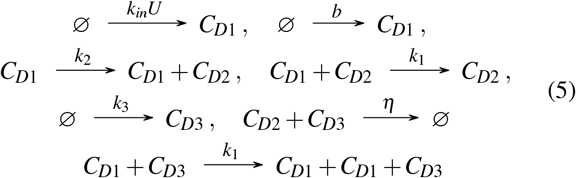

where *k_in_*, *b*, *k*_2_, *k*_1_, *η* ∈ ℝ_+_. Note also that the rate of the degradation process with respect to *C*_*D*1_ considered in [19] is assumed to be zero here.

The dynamics of CRN (5) can be modelled as:

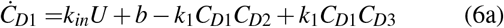

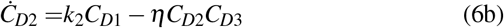

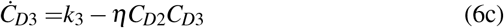

As shown in [19], for any non-negative constant input *U* ^∗^, we obtain a positive locally exponentially stable steady state 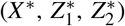.

Through (Jacobian) linearization of system (6), we have for the local dynamics of *BioSD-III*:

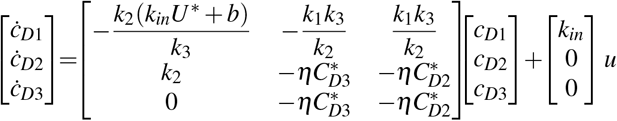

where variables *u* = *U* − *U* ^∗^, 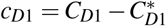, 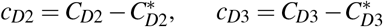 represent small perturbations around 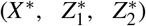. The corresponding input/output relation in the Laplace domain can be described by the following transfer function:

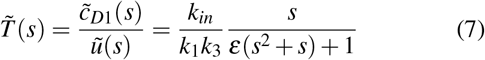

where:

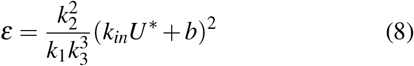

and *s* is the Laplace variable (complex frequency).

Equation (7) is an ideal signal differentiator multiplied by a constant gain in series with a second-order low pass filter. The filtering action can be adjusted to meet our performance standards by appropriately tuning the dimensionless parameter (8). Thus, moving to the time domain, for a given value of (8), there are sufficiently slow input signals yielding:

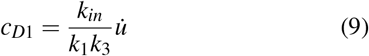

As also shown in [19], the structural complexity of the differentiator module can be reduced by removing the reaction 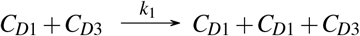 in CRN (5) while the input/output behaviour remains the same. This results in ODE model (6) without the term +*k*_1_*C*_*D*1_*C*_*D*3_ in Equation (6a). However, this simplification, which leads to a different signal differentiator topology called *BioSD-II*, comes with the cost of imposing the following constraint regarding the parameters involved:

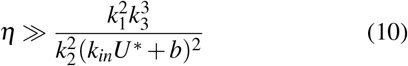

## III. Structure and behaviour of the PID architecture

In this section we present the biological structure of the PID controller this paper focuses on.

In Figure 3 we illustrate the biomolecular architecture of our PID controller regulating a target (output) species, *Y*, of an abstract “cloud” network. The reactions that form the corresponding CRN are:

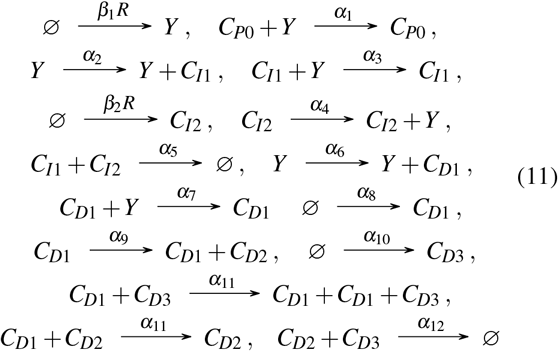

where *α_i_* ∈ ℝ_+_ with *i* ∈ ℕ and 1 ≤ *i* ≤ 12. *R* is a non-negative reference signal that can vary over time and can be controlled externally while *β*_1_, *β*_2_ are non-negative scaling parameters. Through that signal we can adjust the set point of the closed-loop system. In parallel, *C*_*P*0_ can be considered as an auxiliary species with constant concentration that catalyzes the degradation of the target species *Y*. Note also that the modified version of the *antithetic motif* with an additional inhibitory reaction as formed by species *C*_*I*1_, *C*_*I*2_ has been studied in [23].

**Fig. 3:**
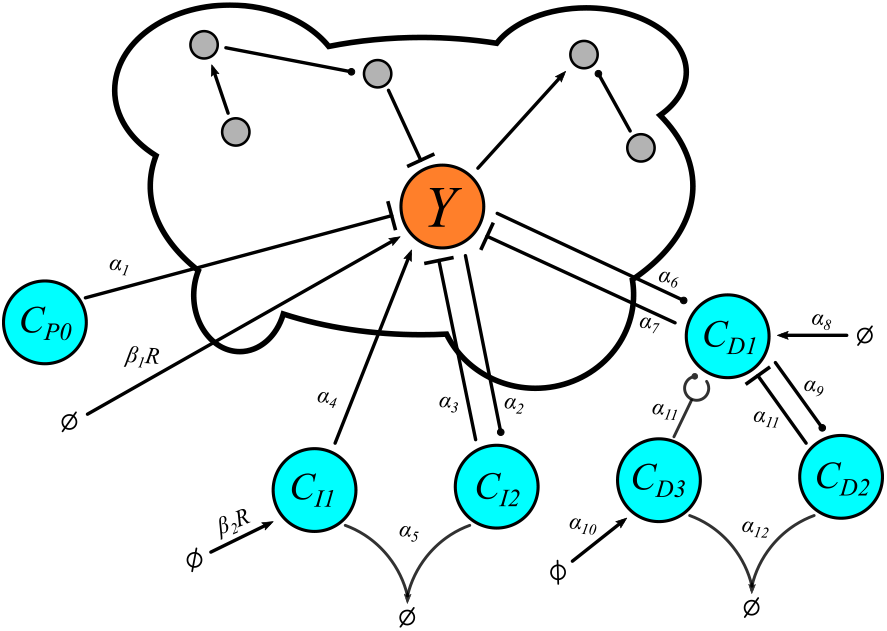
The proposed PID bio-controller regulating a target species which is part of an arbitrary biological process - “cloud” network (CRN (11)).

### A. Achieving PID control

To gain a deeper understanding of the proposed topology (Figure 1B), we study the corresponding dynamics which can be described by the following set of ODEs:

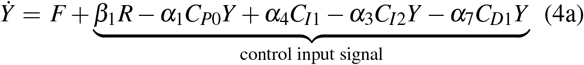

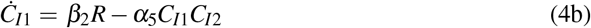

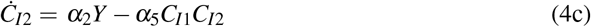

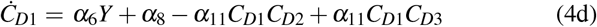

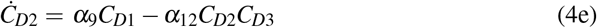

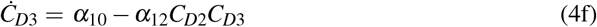

where function *F* represents the mathematical terms describing potential interactions associated with the output species *Y* in the cloud network.

We assume the existence of a (locally) asymptotically stable and biologically meaningful equilibrium for the overall closed loop system for some constant value, *R*^∗^, of reference signal *R* and we focus on the local behaviour of our bio-controller around it. We therefore adopt coordinate transformations of the form *x* = *X* − *X*^∗^ which denote small perturbations around the equilibrium. (*X* and *X* ^∗^ represent any variable involved in the system under consideration and its corresponding steady state, respectively.) Thus, we obtain via (Jacobian) linearization of Equations (4a)-(4f):

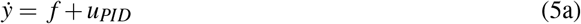

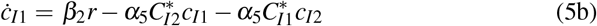

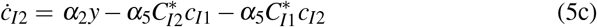

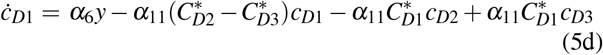

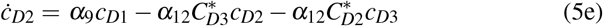

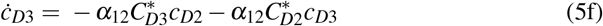

where *f* refers to the “linearized version” of *F* around the aforementioned equilibrium and the control input signal is given by:

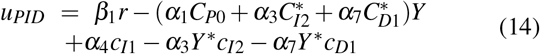

From Equations (4b)-(4c) we get at the steady state :

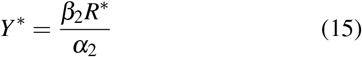

Moreover, species *C*_*D*1_, *C*_*D*2_, *C*_*D*3_ form a *BioSD-III* module with *u* = *y* (see Equations (4d)-(4f)). Thus, taking into account Equations (7)-(10), we have for the input/output relation in the Laplace domain:

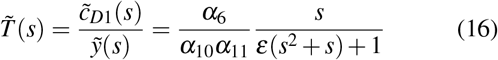

where:

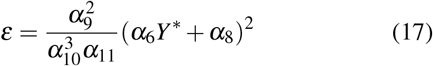

while the parameter constraint for the simplified *BioSD-II* module becomes:

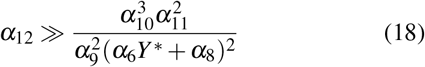

Setting now

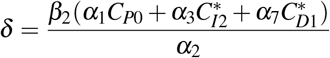

and assuming 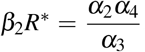, the control input signal (14) can be rewritten as:

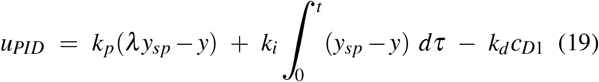

with:

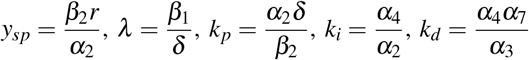

Equation (19) describes a PID control law with *set point weighting* and filtered derivative action. As can be seen, our architecture offers considerable tunability since the controller gains, the set point weight regarding proportional control as well as the filtering action regarding derivative control can be tuned separately as desired. In addition, setting *β*_1_ = 1 or *β*_1_ = 0 leads to a PI-D or I-PD control law, respectively.

## IV. Regulating a specific biological process

In this section we investigate the properties of our PID controller on a specific biological process.

In particular, we replace the abstract cloud network of Figure 3 with a biological process of two mutually activated species, *Y* and *W*, with the first species being the target species on which we apply PID control (Figure 4A). This process is based on a positive feedback loop which is a very common concept in biological systems [24], [25].

**Fig. 4:**
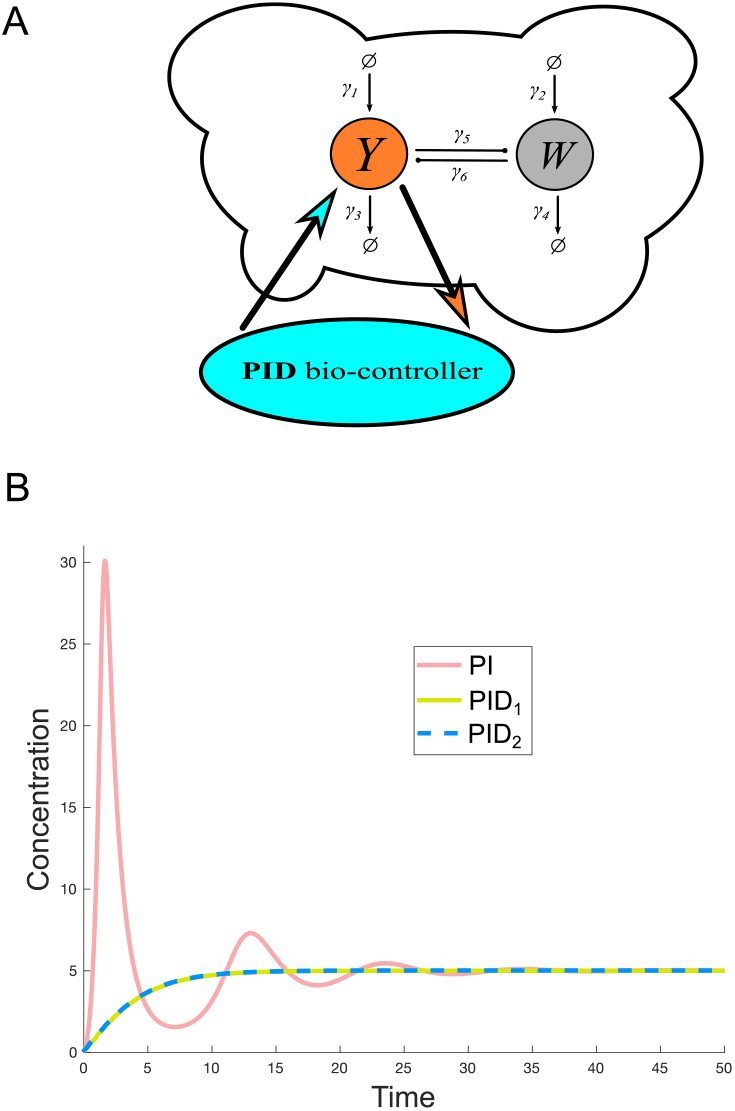
(A) The proposed PID bio-controller regulates a target species (*Y*) of a network consisting of two mutually activated species (CRNs (11), (20)). (B) Simulated response of the target species (*Y*) regarding the topology in (A) described by ODE model (11). In particular: For PI case, we use only Equations (11a)-(11d) with the following parameter values: *β*_2_*R* = 5, *α*_1_*C_P_*_0_ = 0.2, *α*_2_ = 1, *α*_3_ = 0.4, *α*_4_ = 2, *α*_5_ = 10. Additionally, the term −*α*_7_*C*_*D*1_*Y* in Equation (11a) is assumed to be removed since there is no derivative action. In PID_1_ case, derivative control takes place through *BioSD-III*. Here we use Equations (11a)-(11g) with the following parameter values: *α*_6_ = 100, *α*_7_ = 0.15, *α*_8_ = 100, *α*_9_ = 1, *α*_10_ = 100, *α*_11_ = 1, *α*_12_ = 10 while the rest of the parameter values are the same as in PI case. In PID_2_ case we replace *BioSD-III* with *BioSD-II* which results in ODE model (11) without the term +*α*_11_*C*_*D*1_*C*_*D*3_ in Equation (11e). We also use the same parameter values as in PID_1_ case except for *α*_12_ = 500 so that condition (18) is satisfied. The simulations depicted in this figure were performed in MATLAB (Mathworks).

The open-loop process under consideration consists of the following reactions:

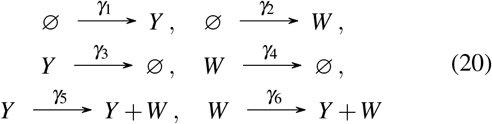

Taking into account CRNs (11) and (20), the dynamics of the resulting closed-loop system can be modelled as:

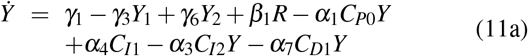

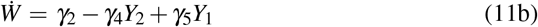

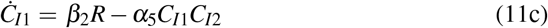

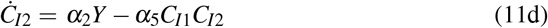

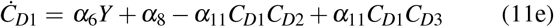

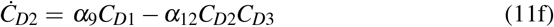

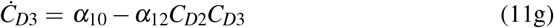

In Figure 4B we present the response of output species, *Y*, using PI and PID control, respectively. In both scenaria identical integral action takes place and, thus, *Y* converges to the same value over time. Nevertheless, the transient response in the first case shows a significant overshoot and oscillations which are eliminated due to the anticipatory action of derivative control in the second case. Moreover, as can be seen, the output response remains the same regardless of the signal differentiator module used in the PID bio-controller.

## V. Stochastic Simulations

The random nature of biomolecular reactions makes biological systems inherently stochastic [18], [27], [28]. The deterministic approach we have followed so far can offer a satisfactory insight into the average biological behaviour when biomolecular populations are sufficiently large. However, this may not be always the case and, as a consequence, analysis of the probabilistic effects may be needed. To this end, we focus here on the stochastic evolution of the closed-loop system shown in Figure 4 over time using the Linear Noise Approximation (LNA). More specifically, in Figure 5 we plot the time evolution of the standard deviation, denoted here as *σ*, with respect to the output species for both PI and PID control. As can be seen, PID control leads to a considerably smaller *σ* compared to PI control at the steady state, demonstrating the noise reduction capability of derivative control through BioSD modules.

**Fig. 5:**
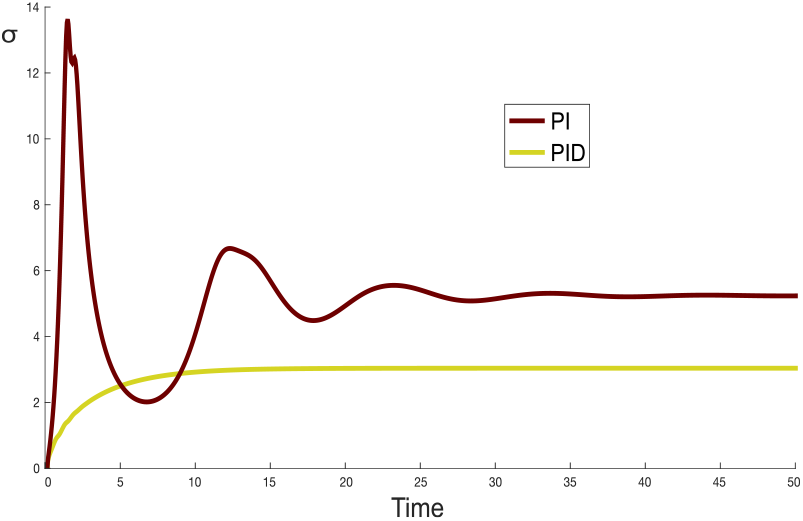
Time evolution of the standard deviation, *σ* of the target species *Y* of the closed-loop system shown in Figure 4. The models and the parameter sets for PI and PID cases correspond to the cases of PI and PID_1_ of Figure 4. Also, the case of PID_2_ results in identical behaviour to PID_1_. The simulations depicted in this figure were performed in Kaemika [26] using LNA.

## VI. CONCLUSION

In this paper we propose a highly tunable CRN architecture capable of applying PID feedback control locally using *set point weights* and derivative control filtering. Notable characteristics of our design are the “antithetic integration” and “BioSD signal differentiation”. About the latter, we consider two differentiator modules of different structural complexity but identical input/output behaviour - reduction of complexity entails addition of a parameter constraint. As far as proportional control is concerned, it is realized through a special birth-death process to which the integral and derivative parts also contribute. To demonstrate the performance benefits of our PID control strategy, we apply it to an (open-loop) process of two mutually activated species and compare it to PI regulation. We show through deterministic simulations that the concentration of the output species of interest exhibits a significantly improved transient response with PID compared to PI control. At the same time, using LNA we show that the addition of *BioSD* derivative action can reduce the standard deviation at the steady state.

An interesting future endeavour would be to study the stochastic behaviour of the proposed controller using other, more accurate methods [29], and compare the results with the LNA ones here. Finally, as our PID bio-controller is experimentally realizable, another interesting future direction would be an *in vitro* implementation via molecular programming. In particular, our topology relies purely on mass action kinetics and, thus, is possible to be translated into a DNA strand displacement system [30], [31], [32].

